# *Drosophila* IMP regulates Kuzbanian to control the timing of Notch signalling in the follicle cells

**DOI:** 10.1101/346585

**Authors:** Weronika Fic, Celia Faria, Daniel St Johnston

## Abstract

The timing of *Drosophila* egg chamber development is controlled by a germline Delta signal that activates Notch in the follicle cells to induce them to cease proliferation and differentiate. Here we report that follicle cells lacking the RNA-binding protein IMP go through one extra division due to a delay in the Delta-dependent S2 cleavage of Notch. The timing of Notch activation has previously been shown to be controlled by cis-inhibition by Delta in the follicle cells, which is relieved when the miRNA pathway represses Delta expression. *imp* mutants are epistatic to *Delta* mutants and give an additive phenotype with *belle* and *dicer* mutants, indicating that IMP functions independently of both cis-inhibition and the miRNA pathway. We find that the *imp* phenotype is rescued by over-expression of Kuzbanian, the metalloprotease that mediates the Notch S2 cleavage. Furthermore, Kuzbanian is not enriched at the apical membrane in *imp* mutants, accumulating instead in late endosomes. Thus, IMP regulates Notch signalling by controlling the localisation of Kuzbanian to the apical domain, where Notch cleavage occurs, revealing a novel regulatory step in the Notch pathway.

**Summary:** IMP regulates Notch signalling in follicle cells by controlling Kuzbanian localisation to the apical domain, where Notch cleavage occurs, revealing a novel regulatory step in the Notch pathway.

## Introduction

RNA binding proteins (RBPs) play diverse roles in the post-transcriptional regulation of gene expression by controlling the splicing, stability, translation or subcellular localisation of specific mRNAs. One of the best studied classes of RBPs is the conserved family of IGF2 mRNA binding proteins (IMPs, also known as the VICKZ family), which are characterised by 4 conserved KH domains, with KH3 and KH4 being most important for RNA-binding, and two N-terminal RRM domains (Degrauwe et al., 2016). Initial studies on IMPs pointed to an important role in mRNA localisation. The *Xenopus* IMP3 orthologue, Vg1RBP/Vera, binds to the localization signal in Vg1 mRNA and co-localises with it to the vegetal cortex of the *Xenopus* oocyte (Deshler et al., 1997; Havin et al., 1998). Similarly, the chicken IMP1, ZBP1 binds to the 54 nucleotide localisation signal in β-actin mRNA to mediate its localisation to the periphery of fibroblasts and the dendrites of neurons (Farina et al., 2003; Tiruchinapalli et al., 2003). However, IMPs also regulate mRNA translation and mRNA stability. Mammalian IMP1-3 were initially identified as translational regulators of Insulin growth factor–like II (IGF-II) mRNA and ZBP1 represses the translation of β-actin mRNA until it reaches its destination (Huttelmaier et al., 2005; Nielsen et al., 1999; Yao et al., 2006). One mechanism by which IMPs regulate mRNA translation and stability is by preventing the binding of siRNAs and miRNAs to their targets, either by masking the binding sites or by sequestering the mRNA away from the Argonaute/RISC complex (Degrauwe et al., 2016). In many cases, IMPs have been found to play an important regulatory role, although the relevant RNA targets have not been identified. For example, IMP1 and 3 are up-regulated in a number of tumours, with their expression levels correlating with increased metastasis and poor prognosis (Ioannidis et al., 2001; Nielsen et al., 2000; Degrauwe et al., 2016).

Vertebrates contain 3 closely-related IMP paralogues, which has hampered functional analysis, whereas *Drosophila* contains a single IMP orthologue with 4 well-conserved KH domains, allowing the genetic analysis of IMP function (Nielsen et al., 2000). IMP was found to bind directly to *oskar* and *gurken* mRNAs and localise with them to the posterior and dorsal sides of the oocyte respectively (Munro et al., 2006; Geng and Macdonald, 2006). Although, the IMP binding sites are required for *oskar* mRNA translation and anchoring, loss of IMP has no obvious phenotype, suggesting that it functions redundantly with other proteins in the germ line. IMP is strongly expressed in the developing nervous system and RNAi knockdown causes neuronal loss and axon pathfinding defects and a reduced number of boutons at the neuromuscular junctions (Boylan et al., 2008; Koizumi et al., 2007). *imp* mutant clones in the developing adult brain cause similar defects in axon elongation in mushroom body neurons, at least in part through IMP’s role in regulating the localisation of *chic* mRNA (Medioni et al., 2014). These neural phenotypes may be related to IMP’s function as temporal identify factor that acts in opposition to Syncrip to specify early-born neuronal fates and to promote neuroblast proliferative capacity (Narbonne-Reveau et al., 2016; Liu et al., 2015). IMP also acts as part of a temporal programme that controls the aging of the testis hub cells. IMP protects *unpaired* mRNA from repression by miRNAs in these cells and as IMP levels fall with age, Unpaired signalling to maintain the male germline stem cells declines, leading to stem cell loss (Toledano et al., 2012).

Here we analyse the function of IMP during the development of the somatic follicle cells of the *Drosophila* ovary and show that it also controls the temporal programme of development in this tissue. Unlike other well-characterised roles of IMP, we find that IMP functions independently of the microRNA pathway to regulate the timing of Delta/Notch signalling.

## Results

### IMP is required for proper timing of Notch signalling in follicle cells

To investigate the role of IMP in the follicle cell layer, we generated clones that were homozygous for the null allele, *imp*^7^, marked by the loss of RFP (Munro et al., 2006). The mutant cells showed no phenotypes during early oogenesis up until the end of stage 6. However, Phalloidin staining of actin revealed that mutant cells at later stages were smaller in size and were more densely packed than the surrounding wild type cells (Fig. 1A’). We observed the same phenotype with a second null allele, *imp*^8^ (Munro et al., 2006). *imp* mutant cells also have smaller nuclei (Fig. 1A, C, C’). The size and number of follicle cells is determined by the timing of the mitotic to endocycle transition, which takes place at stage 6, when the germ cells in the egg chamber produce the DSL ligand, Delta, to activate the Notch pathway in the follicle cells (Deng et al., 2001; Lopez-Schier and St Johnston, 2001). Analysis of 56 *imp* mutant clones revealed that there are twice as many mutant cells in each clone than there are wild type cells in the twin spot clone induced at the same time (Fig. 1B). Thus, *imp* mutant cells go through one extra round of mitosis, suggesting that Delta/Notch signalling is delayed.

**Fig. 1.**
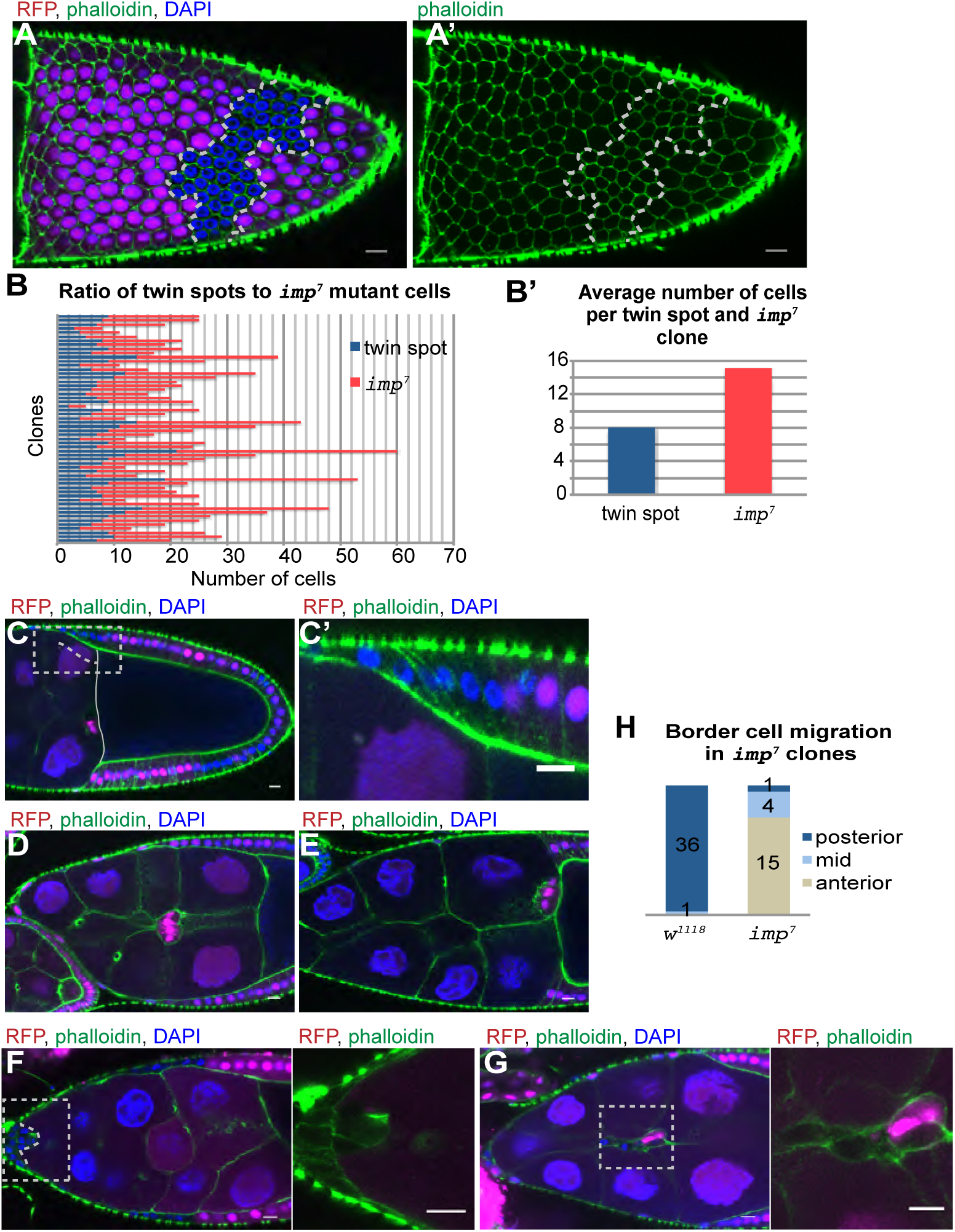
*imp* mutant cells go through one extra division. (A) Surface view of a stage 10a egg chamber containing an *imp^7^* mutant follicle cell clone (marked by the loss of RFP, magenta) stained with phalloidin (green) and DAPI (blue). The mutant cells are outlined in white. (B) A graph showing the number of wild-type and *imp^7^* mutant cells in 56 independent twin spot clones. (B’) A histogram showing the average number of cells per wild-type clone (n=8) and *imp^7^* clone (n=16, B’). (C,C’) Stage 10a chamber with an *imp^7^* follicle cell clone that has not yet migrated posteriorly to envelop the oocyte. (D,E) Wild-type stage 9 (D) and stage 10b (E) egg chambers showing the migration of the border cells between the nurse cells to reach the anterior of the oocyte at stage 10b. (F) A stage 9 egg chamber with an *imp^7^* mutant clone that includes all of the border cells, which have failed to detach from the anterior. (G) A stage 9 egg chamber containing a mosaic of *imp^7^* mutant and wild-type border cells. The mutant cells are found at the back of the cluster and dragged by leading wild type cells. The migration of these clusters is severely delayed and they are often move only half way to the oocyte. (H) Quantification of how far wild-type border cell clusters and entirely mutant clusters have moved by stage 10b.

Notch activation controls both the mitosis to endocycle switch and follicle cell differentiation. The late differentiation in the absence of IMP leads to delays in several other aspects of follicle cell development. For example, the lateral follicle cells move posteriorly to form a columnar epithelium around the oocyte during stages 8-9, but *imp* mutant cells do this more slowly and later than normal (Fig. 1C, C’). During stage 9 of oogenesis, the anterior most follicle cells adopt the border cell fate, delaminate from the epithelium and migrate between the nurse cells to the anterior of the oocyte (Fig. 1D, E). When the entire border cell cluster is mutant for *imp*, the cells frequently fail to delaminate and remain at the anterior of the egg chamber, while those that do delaminate often only migrate part of the way to the oocyte (Fig. 1F, H). When the cluster contains both mutant and wild-type cells, the wild-type cells lead the migration with the mutant cells trailing behind (Fig. 1G). Thus, loss of IMP affects the timing of all aspects of follicle cell behaviour suggesting that it plays a general role in this process. The localisation of IMP does not give any clues as it its function, however, as IMP protein is uniformly distributed throughout the cytoplasm of the follicle cells (Fig. S1).

To test whether IMP is required for the proper timing of Notch pathway activation in the follicle cells, we stained *imp* mutant clones for Cut and Hindsight (Hnt). Cut is expressed from stages 1 to 6 of oogenesis and is down-regulated at stage 7 in response to Notch activation (Sun and Deng, 2005). By contrast, Hnt is expressed only in post-mitotic cells that already received the Delta signal from the germ line (Sun and Deng, 2007). *imp* clones continue to express Cut at stage 7 in contrast to wild type cells in the same egg chamber (Fig. 2A). However, Cut expression is lost in the majority of mutant cells at stage 8 (Fig. 2C). On the other hand, Hnt is not expressed in *imp* mutant cells at stage 7 as in wild-type, but appears one stage later (Fig. 2B, D). Notch activation therefore occurs later than normal in *imp* mutant follicle cells, resulting in the delay of follicle cell differentiation.

**Fig. 2.**
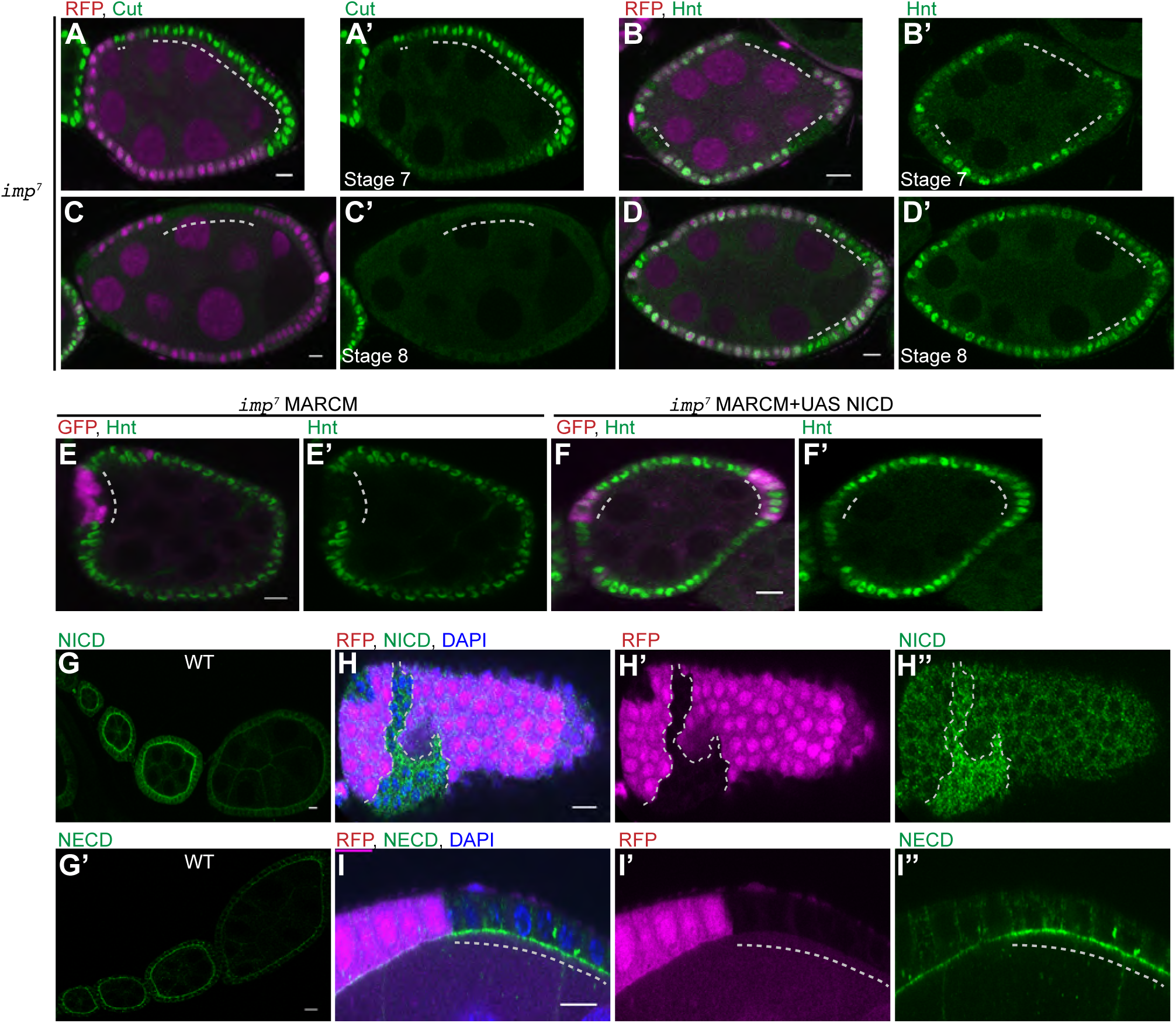
IMP is required for the 1^st^ cleavage of Notch. (A, A’) Cut expression in a stage 7 egg chamber containing an *imp^7^* follicle cell clone marked by the loss of RFP. Cut is only expressed in the mutant cells. (B,B’) Hnt expression in a stage 7 egg chamber containing several *imp^7^* follicle cell clones marked by the loss of RFP. Hnt is expressed in the wild-type cells, but not in the mutant cells at stage 7. (C-D’) Stage 8 egg chambers containing *imp^7^* follicle cell clones stained for Cut (C,C’) and Hnt (D,D’). The mutant cells no longer express Cut and have turned on Hnt at this stage. (E-F’) Stage 7 egg chambers containing *imp^7^* MARCM clones marked by the expression of GFP (magenta) and stained for Hnt (green). The mutant cells do not express Hnt at stage 7, but this is rescued by the expression of the Notch intracellular domain, NICD. (G,G’) Expression of the Notch intracellular domain (NICD, G) and the Notch extracellular domain (NECD, G’) in early stage egg chambers. Both NICD and NECD are enriched at the apical side of the follicle cells until stage 6 when they are down-regulated as a result of Delta signaling. (H-I’) NICD (H-H”) and NECD (I-I”) are not down-regulated in *imp^7^* mutant cells (marked by the loss of RFP).

To directly test whether impaired Notch signalling is responsible for the *imp* phenotype, we used the MARCM system to express a constitutively-active form of Notch, the Notch intracellular domain (NICD), in *imp* mutant cells (Lee and Luo, 2001; Go et al., 1998). Control *imp* mutant MARCM clones do not turn on Hnt at stage 7, but expressing NICD in the mutant cells restores timely Hnt expression (Fig. 2E, F). This indicates that IMP controls Notch activity upstream of NICD production.

### IMP acts at or before the first cleavage of Notch

Upon binding to its ligand, Delta, the extracellular domain of Notch is cleaved at S2 site by the ADAM10 protease, Kuzbanian, to produce a transient form of the receptor, NEXT, which contains the transmembrane and intracellular domains of Notch (Pan and Rubin, 1997; Lieber et al., 2002). NEXT then undergoes a second cleavage at the S3 site mediated by the Presenilin/γ secretase complex (De Strooper et al., 1999; Struhl and Greenwald, 1999; Vaccari et al., 2008; Ye et al., 1999). This releases the intracellular domain of Notch (NICD), which translocates to the nucleus to regulate transcription in association with Suppressor of Hairless protein (Su(H)) (Bray, 2016).

Stainings with antibodies that detect the Notch intracellular and extracellular domains reveal that full-length Notch accumulates on the apical side of the follicle cells during early oogenesis and is then cleared from the membrane at stage 6, when signalling occurs (Fig. 2G, G’). However, both antibodies detect high levels of Notch at stage 7-8 in *imp* mutant cells (Fig. 2H, I). The Notch extracellular domain is removed by the first cleavage and then endocytosed with Delta into the signalling cell (Parks et al., 2000; Nichols et al., 2007; Langridge and Struhl, 2017). The persistence of the extracellular domain signal in the mutant cells therefore indicates that loss of IMP inhibits Notch signalling before or at the first, Kuzbanian-dependent cleavage of the Notch extracellular domain.

To test whether IMP is more generally involved in Notch signalling, we generated mutant clones in the wing imaginal disc. Notch activity is required for Cut expression in two rows of cells along the dorsal-ventral midline of the wing disc (Fig. 3A), (Micchelli et al., 1997). Large *imp* mutant clones that include the dorsal-ventral boundary have no effect on Cut expression, however (Fig. 3B). Furthermore, adult wings containing *imp* clones have a normal bristle pattern and never show any of the wing–notching characteristic of *Notch* mutants, although they have some wing venation defects. *imp* null mutant clones in the eye imaginal disc also showed no phenotype (Fig. 3C). Thus, IMP seems to be specifically required for Notch activation in the ovary and is not a general component of the Notch signalling pathway.

**Fig. 3.**
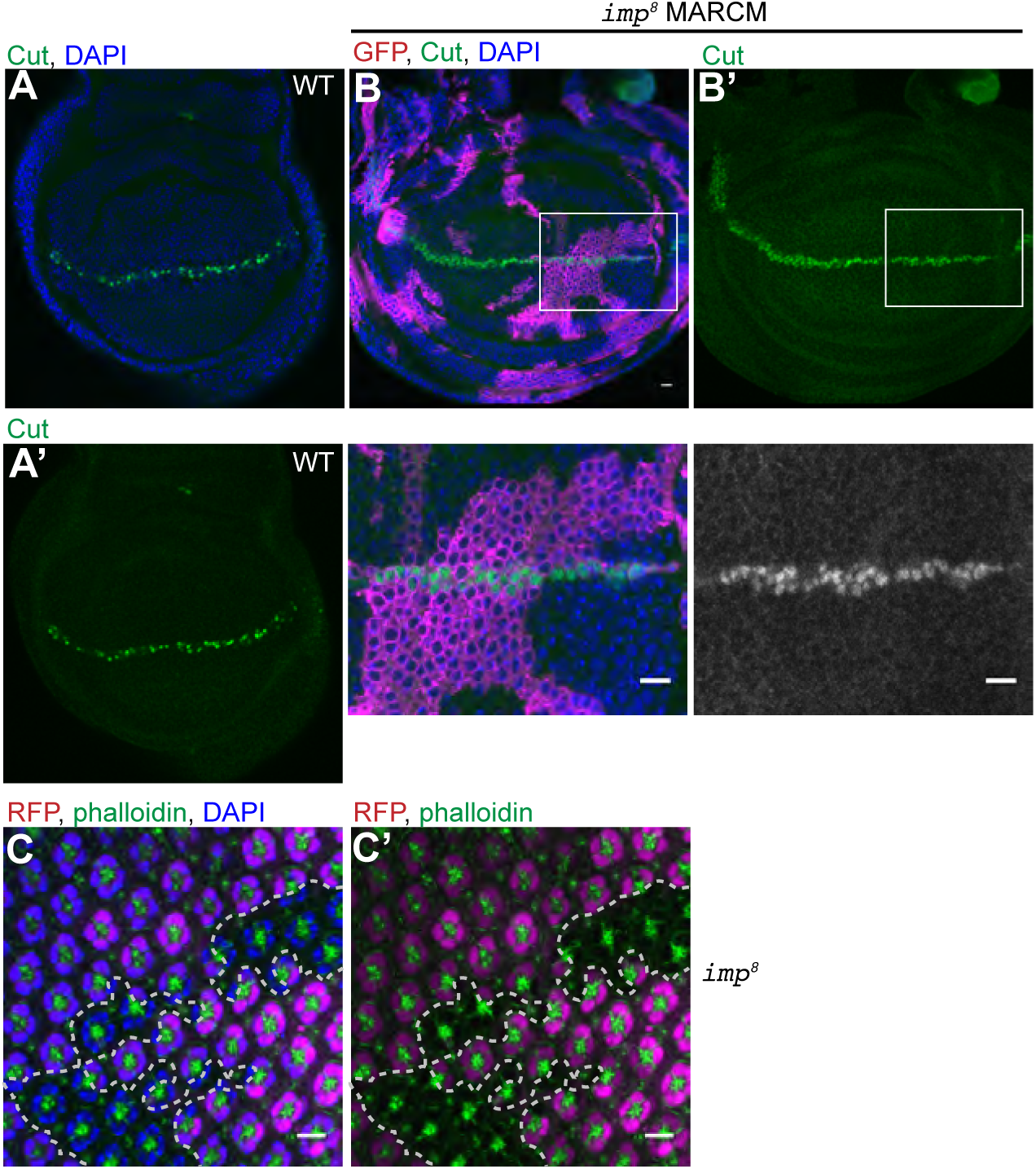
IMP is not a general component of Notch signalling pathway. (A-B) 3^rd^ larval instar wing imaginal discs stained for Cut (green). (B) shows a disc containing *imp^8^* MARCM clones marked by GFP expression (magenta). Cut is expressed normally along the dorsal-ventral compartment boundary in mutant clones. (C) A 3^rd^ instar eye imaginal disc containing *imp^8^* mutant clones marked by the loss of RFP. The mutant cells are indistinguishable from wild-type.

### *imp* does not act through the micro RNA pathway

In addition to binding to Notch in trans to activate its cleavage and signalling, Delta expressed in the same cell can bind to Notch in cis to inhibit signalling (de Celis and Bray, 1997; Micchelli et al., 1997; Cordle et al., 2008; Sakamoto et al., 2002; Miller et al., 2009). Indeed cis-inhibition by Delta expressed in the follicle cells controls their competence to respond to Delta from the germ line, as *Delta* mutant follicle cell clones switch from mitosis to the endocycle too early and undergo precocious differentiation (Poulton et al., 2011). This inhibition is regulated by the microRNA pathway, which represses Delta expression in the follicle cells to relieve cis-inhibition at stage 6. Mutants in the conserved components of the microRNA pathway, *belle* and *dicer-1* therefore cause a similar delay in follicle cell development to *imp* mutants (Poulton et al., 2011). As IMP is an RNA-binding protein that modulates the miRNA pathway in other contexts (Toledano et al., 2012), this raises the possibility that it is required for the miRNA-dependent repression of Delta in the follicle cells.

We compared the phenotypes of *belle* and *dicer-1* mutants with that of *imp* to confirm that they cause a similar inhibition of Notch activation. Like *imp* mutants, *belle* null mutant follicle cells go through one extra division and border cell migration is disrupted (Fig. 4A, B, C). Furthermore, *belle* and *dicer-1* mutant cells accumulate uncleaved Notch at their apical surfaces, as shown by the persistence of staining with antibodies against the Notch extra-cellular and intra-cellular domains (Fig. 4D, E; Fig. 2A, B). If IMP functions in the same micro RNA pathway as Belle and Dicer-1, double mutants should show an identical delay in follicle cell differentiation as *imp*, *belle* and *dicer-1* single mutants, as the removal of a second essential component of the pathway should have no further effect. On the other hand, one would expect an additive effect if IMP functions in a parallel pathway. We therefore generated *imp* mutant clones marked by the loss of RFP and *belle* or *dicer-1* mutant clones marked by the loss of GFP in the same egg chambers. This allowed us to directly compare the phenotypes of cells mutant for either gene with the double mutant cells that express neither GFP or RFP. While the single mutant clones produced the expected phenotypes, the double mutant cells showed a more severe delay in Notch activation, as illustrated by their continued expression of Cut after the single mutant cells have lost expression (Fig. 4F and Fig. 2C). This suggests that IMP functions in a different pathway from Belle and Dicer-1 to control the timing of Notch activation in the follicle cells.

**Fig. 4.**
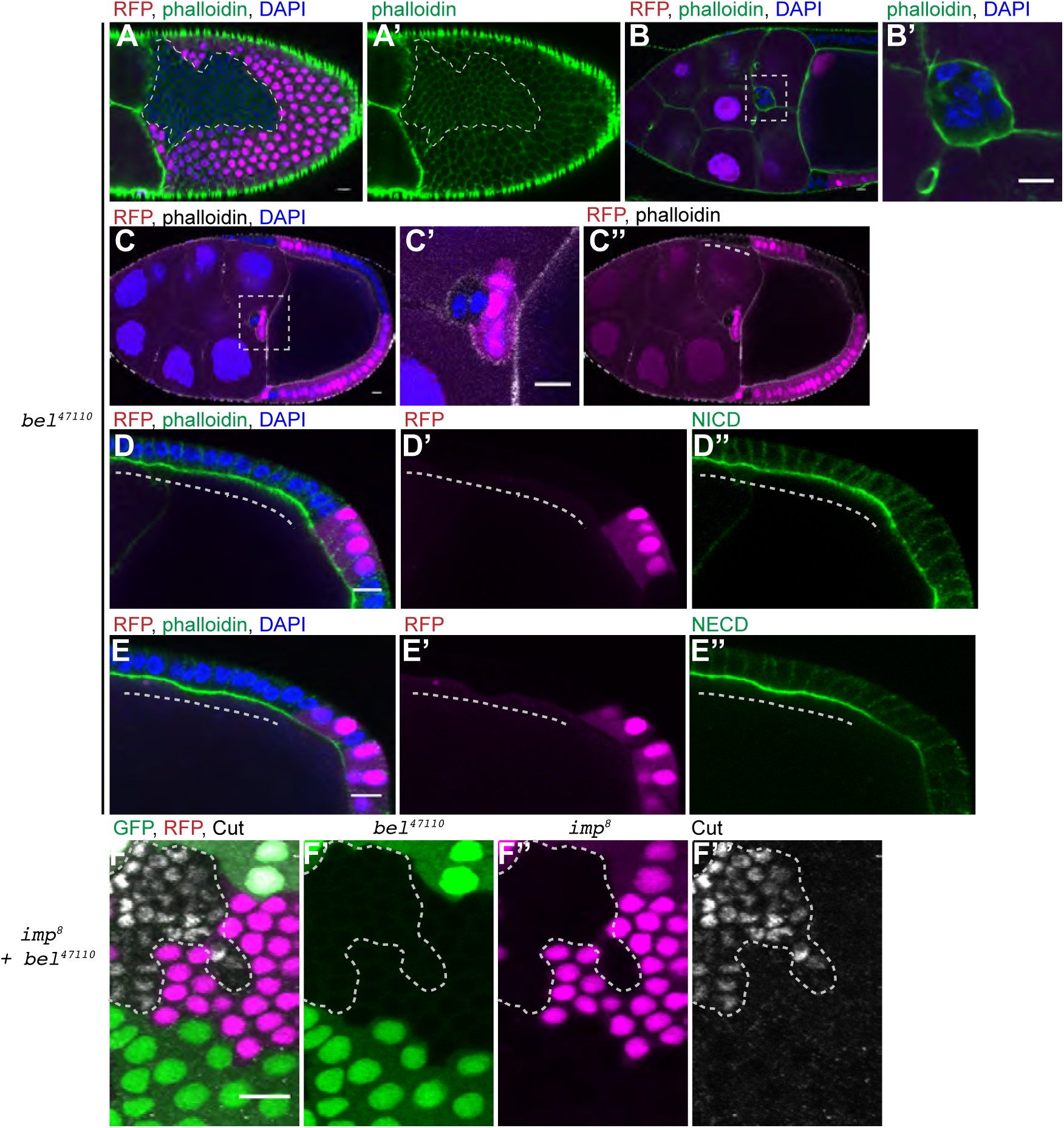
IMP does not act through the micro RNA pathway. (A-C) Stage 10a egg chambers containing *belle*^47110^ mutant clones marked by the loss of RFP. The mutant cells show a similar phenotype to *imp*. They go through one extra round of division and are therefore smaller than the wild-type cells (A). When all of the border cells are mutant, there is a delay in border cell migration (B). In mosaic border cell clusters, the mutant cells lag behind the wild-type border cells (C-C”). The migration of mutant follicle cells to envelop the oocyte is also delayed (white dashed line in C’). (D-E) Stage 9 egg chambers *belle*^47110^ mutant clones marked by the loss of RFP stained for NICD and NECD. The mutant cells retain high levels of NICD (D-D”) and NECD (E-E”) at their apical membranes. (F-F”) A stage 9 egg chamber containing both *imp^7^* mutant clones marked by the loss of RFP (magenta) and *belle*^47110^ mutant clones marked by the loss of GFP (green), stained for Cut (white). Cut is still expressed in the double mutant cells (marked by the dashed line), but not in the single mutant cells.

### *imp* is epistatic to *Delta*

Although IMP does not appear to function in the micro RNA pathway, it could still control Notch activation by repressing Delta expression in the follicle cells to relieve cis-inhibition. If this is the case, *Delta* should be epistatic to *imp*, with double mutant clones showing the *Delta* phenotype of early differentiation and exit from the cell cycle. We first confirmed that *Delta* clones cause premature differentiation of the follicle cells and observed that mutant cells have larger nuclei, indicating premature entry into endocycle (Fig. 5A). Furthermore, *Delta* mutant cells turn off Cut expression before stage 6, unlike wild-type cells, indicating that the cells lacking Delta differentiate precociously (Fig. 5B).

To examine the epistatic relationship between *Delta* and *imp*, we generated clones of null mutants in each gene in the same egg chambers, marked by the loss of GFP and RFP respectively. This revealed that *Delta*, *imp* double-mutant cells show the same delay in Notch signalling as *imp* single mutant cells (Fig. 5C, D, E). Double mutant clones express Cut protein at stage 6/7, as do neighbouring wild type cells, whereas *Delta* single mutant clones have already turned Cut off at this stage (Fig. 5D). Moreover, Hnt expression is not switched on prematurely in *Delta*, *imp* clones, as it is in *Delta* clones (Fig. 5C). More importantly, like single *imp* mutant clones, the *Delta imp* double mutant clones express Cut longer than wild-type cells, (Fig. 5E). To confirm this result, we also examined whether IMP regulates *Delta* mRNA stability or translation, using a sensor line that contains the 3UTR of *Delta* downstream of the GFP coding sequence (Poulton et al., 2011). However, *imp* clones expressed the same level of GFP from the *Delta* sensor as wild-type cells (Fig. 3A). The observation that the *imp* mutants are epistatic to *Delta* mutants in the follicle cells indicates that IMP regulates the timing of Notch activation independently of and in parallel to Delta cis-inhibition.

**Fig. 5.**
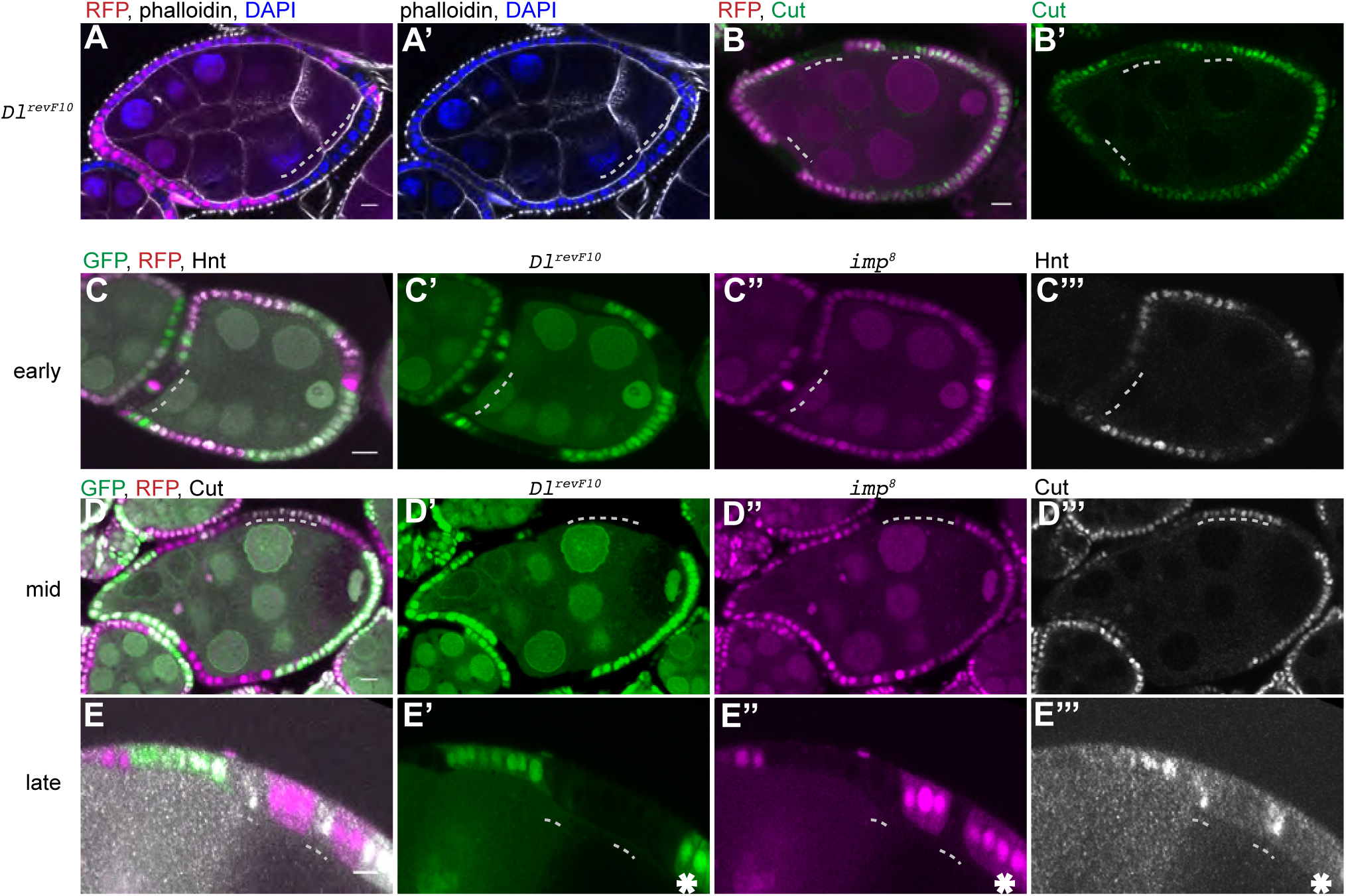
*imp* is epistatic to *Delta* in follicle cells. (A, B) Stage 6 egg chambers containing *Dl*^revF10^ mutant cells marked by the loss of RFP. The mutant cells are larger than wild-type and have bigger nuclei, indicating that they have undergone the switch from mitosis to endoreplication early (A,A’). The *Dl* mutant cells switch off Cut expression (green) before the wild-type cells (B, B’). (C-E) Egg chambers containing both *Dl*^revF10^ clones marked by the loss of GFP (green) and *imp^8^* mutant clones marked by the loss of RFP (magenta). (C-C”’) *imp Dl* double mutant cells do not express Hnt (white) at stage 6/7, whereas *Dl* mutant cells do. (D-D”’) *imp Dl* double mutant cells still express Cut (white) at stage 6, unlike *Dl* mutant cells. (E-E”’) *imp Dl* double mutant cells (marked by the dashed lines) still express Cut at stage 7, in contrast to wild-type cells (marked with an asterisk).

### Kuzbanian is indispensable for Notch pathway activation in follicle cells

Since IMP is only required in the follicle cells, it cannot affect the expression of Delta in the germ line and must therefore affect either the ability of Notch to bind to Delta in trans or the first Delta-dependent cleavage of Notch at the S2 site, which is mediated by the ADAM family metalloprotease, Kuzbanian (Pan and Rubin, 1997; Lieber et al., 2002). To confirm that Kuzbanian is required for Notch signalling in the follicle cells, we generated clones of an amorphic allele, *kuz*^e29-4^ (Rooke et al., 1996). *kuz* mutant follicle cells show an identical phenotype to null mutations in Notch: mutant cells continue dividing after stage 6, leading to an increase in cell number and a decrease in cell size, and fail to differentiate, as they continue to express Cut and never turn on Hnt (Fig. 6A, B). Staining for the extra-cellular and intra-cellular domains of Notch showed that *kuz* mutant cells maintain high levels of NICD and NECD at the follicle cell apical membrane, consistent with its role in the first cleavage of Notch, a phenotype that resembles that of *imp* (Fig. 6C, D).

**Fig. 6.**
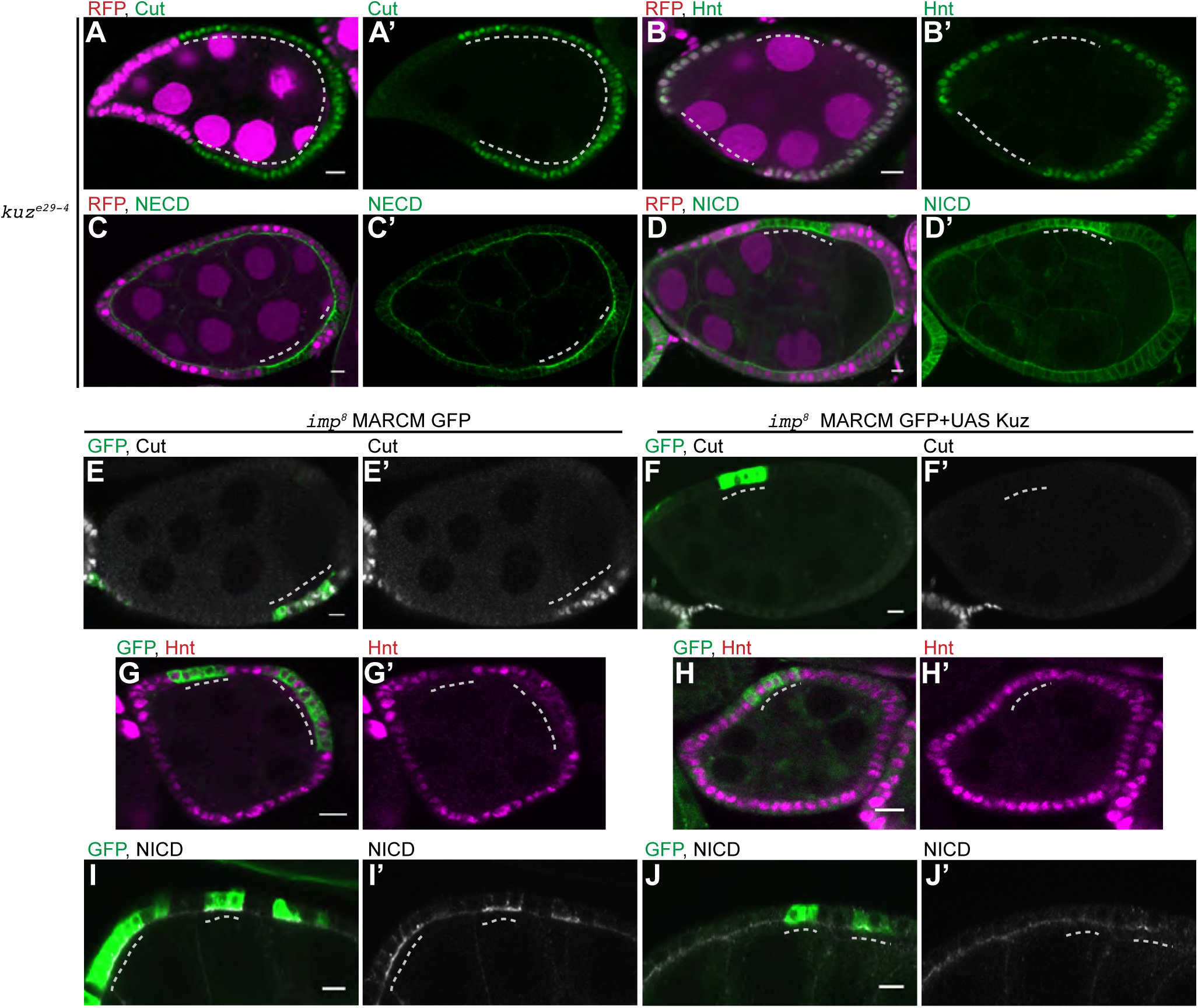
Kuzbanian is required for Notch activation in the follicle cells. (A-D) Egg chambers containing *kuz*^e29-4^ mutant cells marked by the loss of RFP. The *kuz* mutant cells continue to express Cut at stage 7 (A, A’) and do not express Hnt (B, B’). Loss of Kuz leads to the persistence of high levels of NECD (C, C’) and NICD (D, D’) at the apical membrane of the follicle cells after stage 6. (E-J’) *imp*^8^ MARCM clones marked by the expression of GFP (green) without any additional transgenes (E,G,I) or with UAS-Kuz (F,H, J). Kuzbanian expression in *imp* mutant cells restores the timely repression of Cut (F,F’) and activation of Hnt (H,H’), in contrast to control *imp* mutant cells at stage 7 (G, G’& I, I’). Control *imp* mutant clones retain high levels of NICD at the apical membrane of the follicle cells after stage 6 (I, I’), whereas NICD is down-regulated in *imp* mutant cells expressing Kuzbanian, as in wild-type cells (J,J’).

Since *kuz* and *imp* mutants block the same step in Notch activation, we asked whether the *imp* phenotype results from a deficit in Kuzbanian activity by over-expressing Kuzbanian in *imp* mutant cells using the MARCM system (Lee and Luo, 2001). Control *imp* MARCM clones continue to express Cut and not Hnt at stage 7, as expected (Fig. 6E, G). By contrast, expression of Kuz eliminates the delay in Cut repression and restores timely Hnt expression (Fig. 6F, H). Furthermore, while Notch remains at high levels in the apical plasma membrane of *imp* mutant cells at stage 7, it disappears on schedule in mutant cells over-expressing Kuzbanian, as it does in wild-type (Fig. 6I, J). Thus, increasing the levels of Kuzbanian rescues the Notch signalling defect in *imp* mutant follicle cells, suggesting that IMP is required for normal Kuzbanian activity.

### IMP is required for the apical accumulation of Kuzbanian

Because no anti-Kuz antibodies are available, we took advantage of a GFP tagged *kuzbanian* BAC transgene that rescues the lethality of *kuz* mutants (Dornier et al., 2012). Kuz-GFP is expressed at very low levels in the follicle cells, with the highest expression at stages 5-6. Live imaging of egg chambers with two copies of this transgene revealed that Kuz-GFP is enriched at the apical side of the follicle cells and in intracellular speckles (Fig. 7A, A’, A”). This weak apical enrichment is reduced or lost in the majority of *imp* mutant cells (63% of 38 clones, Fig. 7B, B’). Instead, most mutant cells show a marked increase in the number and brightness of the speckles, with 56% of clones containing large intracellular foci (Fig. 7C, D). These structures partially co-localize with CellMask, which stains the plasma membrane and endocytic compartments (Fig. 7E, E’, E”). Unlike the signal at the apical membrane, the intracellular foci of Kuzbanian remain intact after fixation, which allowed us to stain for markers for different vesicular compartments. The Kuzbanian-positive punctae showed the strongest co-localisation with Rab 7, a marker for late endosomes, whereas they did not co-localise with the Golgi marker, GM130 (Fig. 7F, G). These results suggest that loss of IMP disrupts the intracellular trafficking of Kuzabanian, leading to a reduction in its levels at the apical plasma membrane, where Notch cleavage occurs.

**Fig. 7.**
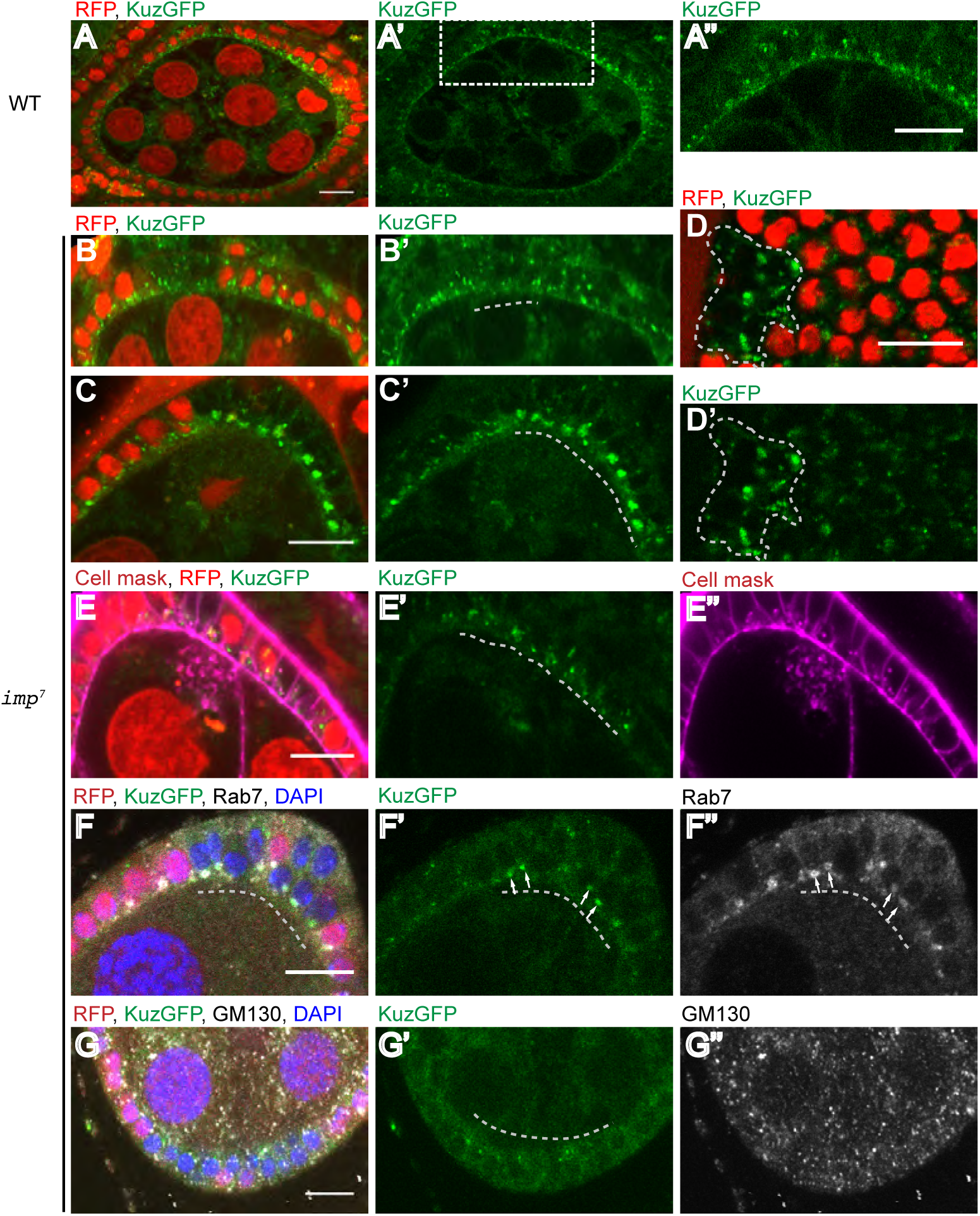
Loss of IMP disrupts Kuzbanian localization. (A-G”) Live egg chambers from females carrying two copies of a Kuz-GFP BAC transgene. (A,A’) Kuzbanian-GFP localises to the apical membrane of the follicle cells and to intra-cellular punctae. (B-D’) *imp*^7^ mutant cells marked by the loss of RFP (red) show a decrease in the amount Kuz-GFP at the apical membrane (B,B’) and an increase in the Kuz-GFP found in bright intracellular foci (C, C’, D, D’). (E, E’, E”) Kuz-GFP intracellular speckles partially co-localise with CellMask. (F-G”) The large intracellular Kuz-GFP foci co-localise with Rab7 (white in F,F”), a marker for late endosomes, but do not co-localise with the Golgi marker, GM130 (white in G,G”).

## Discussion

Although Delta/Notch signalling uses a very simple signal transduction pathway, its activity can be regulated at several levels to control the timing and direction of signalling in context-dependent manner. Factors controlling the endocytosis of Notch, such as Numb, can determine the direction of signalling, whereas the ubiquitin ligases that induce Delta endocytosis, Neuralised and Mindbomb determine where and when signalling occurs (Pavlopoulos et al., 2001; Deblandre et al., 2001; Lai et al., 2001; Itoh et al., 2003; Le Borgne et al., 2005; Lai et al., 2005). Whether cells respond to a particular DSL ligand can also be regulated by the glycosylation of the Notch extracellular domain by the Fringe family of proteins, making *Drosophila* Notch more responsive to Delta and less responsive to Serrate (Moloney et al., 2000; Bruckner et al., 2000; Goto et al., 2001). Indeed, the modification of Notch by Fringe in the polar/stalk follicle cell precursors renders these cells more responsive to Delta than the other follicle cells, thereby restricting polar cell fate to the ends of the early egg chamber (Grammont and Irvine, 2001). Finally, signalling can be modified by cis-inhibition by DSL ligands, which impede Notch interactions with activating ligands in trans. This mechanism plays a role in controlling the timing of follicle cell differentiation until Delta is down-regulated in the follicle cells by the micro RNA pathway (Poulton et al., 2011). Here we present evidence for a new mechanism that regulates the activity of the Notch pathway through the localisation of the ADAM metalloprotease Kuzbanian.

Our results show that mutants in the RNA-binding protein, IMP, have impaired Delta/Notch signalling in mid-oogenesis, which results in one extra follicle cell division. Although mutants in the microRNA pathway give a very similar phenotype to *imp,* because of a failure to repress Delta translation, IMP functions in parallel to this pathway. Firstly, double mutants between *imp* and *belle* or *dicer-1* show an additive delay in follicle cell differentiation. Secondly, *imp* mutants are epistatic to *Delta* mutants, indicating that IMP does not function by relieving cis-inhibition. Instead, we find that the delay in Notch signalling can be efficiently rescued by over-expression of Kuzbanian. Furthermore, loss of IMP disrupts the enrichment of Kuzbanian at the plasma membrane, which is where Notch must be cleaved for signalling to occur. Thus, IMP is required in some way for the localisation of Kuzbanian to the site where germline Delta binds to Notch in trans to trigger the first cleavage, identifying a new regulatory step in the Notch pathway.

The loss of IMP does not disrupt other Delta/Notch signalling processes, such as the formation of the dorsal ventral boundary in the wing, indicating that it is not a general component of the pathway. This raises the question of why IMP is specifically required in the follicle cells. One possibility is that this relates to the different geometry of the Delta/Notch interaction in the follicle cells, compared to other signalling events. Most examples of Delta/Notch signalling occur between adjacent cells in epithelial tissues, where Delta and Notch can only interact at the lateral membrane, usually at the level of the adherens junctions (Woods and Bryant, 1993; Bray, 2016; de Celis et al., 1996; Micchelli and Blair, 1999; Kidd et al., 1989; Andersson et al., 2011). By contrast, the germ cells of the egg chamber, which produce the activating Delta signal, contact the apical side of the follicle cells, and both Notch and Kuzbanian therefore need to be localised apically for signalling to occur. Notch localisation is not affected by *imp* mutants, however, as it accumulates at high levels at the apical side of mutant cells. Thus, IMP appears to specifically disrupt Kuzbanian localisation rather affecting apical trafficking more generally.

Since IMP is a cytoplasmic RNA-binding protein, it presumably acts by regulating the stability, translation or localisation of specific mRNAs. One possibility is that IMP acts on *kuz* mRNA directly. Indeed, we have observed by qPCR that *kuz* RNA is enriched in immuno-precipitations of IMP (data not shown), although we cannot detect the mRNA by fluorescent in situ hybridisation, presumably because it is present at very low levels. It seems unlikely that IMP regulates the translation of *kuz* mRNA, as Kuzbanian protein levels do not appear to change in *imp* mutants. There is also no evidence for a role of IMP in the localisation of *kuz* mRNA, since IMP itself is not localised and Kuzbanian is a secreted transmembrane protein that must be translated at the endoplasmic reticulum and trafficked to the cell surface through the Golgi complex. One possibility is that IMP controls Kuzbanian localisation through 3’UTR-dependent protein localisation in a similar way to that in which HuR binds to the long 3’UTR of CD47 mRNA to recruit SET protein, which then facilitates CD47 protein trafficking to the cell surface (Berkovits and Mayr, 2015). In this scenario, IMP binding to the 3’UTR of *kuz* mRNA would recruit a co-factor that then associates with the cytoplasmic domain of Kuzbanian protein to direct its trafficking to the apical plasma membrane.

It seems more likely that IMP enhances the translation or stability of another mRNA that encodes a factor that either directs the apical trafficking of Kuzbanian or anchors it in the apical plasma membrane. Possible candidates include the TspC8 Tetraspanins, Tsp86D and Tsp3A, which have been shown to enhance the trafficking of Kuzbanian to the cell surface in S2 cells and in the migrating border cells (Dornier et al., 2012). However, the accumulation of Kuzbanian protein in Rab-7 positive late endosomes in *imp* mutant cells suggests that Kuzbanian is being endocytosed from the apical membrane in the absence of IMP, arguing that the phenotype results from the loss of a factor that stabilises Kuzbanian at the membrane and prevents its endocytosis, rather than a factor that facilitates its delivery there. Various cross-linking approaches, such as RIPseq, i-CLIP, PAR-iCLIP and eCLIP have shown that IMPs bind to 100-1000s of mRNAs, predominantly in their 3’UTRs (Conway et al., 2016; Hansen et al., 2015). IMP binding sites are particularly enriched in mRNAs encoding cytoskeletal components and trafficking factors, many of which are plausible candidates for the relevant target of IMP in regulating Kuzbanian localisation and Notch signalling in the follicle cells.

## Materials and methods

### Drosophila mutant stocks and transgenic lines

We used the following mutant alleles and transgenic constructs: *imp^7^* and *imp^8^* (Munro et al., 2006), GFP-IMP trap line 126.1 (personal communication Alain Debec, Morin et al, 2001), UAS-NICD (Go et al., 1998) a gift from S.Bray, University of Cambridge, Cambridge), *bel^47110^* and Dl-3’UTR sensor (Poulton et al., 2011), *Delta^revF10^* and *Delta^M1^*(Sun and Deng, 2005) a gift from S.Bray, University of Cambridge, Cambridge), dcr-1Q^1147X^ (Lee et al., 2004) a gift from A. Brand, The Gurdon Institute, Cambridge), Kuz-GFP and *kuz^e29-4^* (Dornier et al., 2012; Rooke et al., 1996), UAS-kuz (Sotillos et al., 1997; Kyoto Stock Center 108440). The following stocks were used to generate mitotic clones: ubiRFP-nls, hsflp, FRT19A (BDSC 31418), FRT40A ubiRFP-nls (BDSC 34500), FRT82B, ubiRFP-nls (BDSC 30555) and FRT82B ubiGFP (BDSC 5188). Mosaic analysis with a repressible cell marker, MARCM; after the method of (Lee and Luo, 2001) was carried out using UAS-GFP-mCD8 (Lee and Luo, 1999) as the marker.

### Reagents

The following antibodies were used: mouse anti-Cut (Blochlinger et al., 1990, Developmental Studies Hybridoma Bank, 2B10), mouse anti-Hnt (Yip et al., 1997, Developmental Studies Hybridoma Bank, 1G9), mouse anti-NICD (Fehon et al., 1990, Developmental Studies Hybridoma Bank, C17.9C6), mouse anti-NECD (Diederich et al., 1994, Developmental Studies Hybridoma Bank, C458.2H). All primary antibodies from DSHB were used at a dilution of 1:100. Anti-GM130 was purchased from Abcam (Sinka et al., 2008, ab30637) and used at 1:500. Anti-Rab7 (rabbit) was kindly provided by Nakamura lab and used at 1:1000 (Tanaka and Nakamura, 2008). Conjugated secondary antibodies were purchased from Jackson ImmunoResearch and used at a dilution of 1:1000. F-actin was stained with AlexaFluor 568 or AlexaFluor 647 Phalloidin (Invitrogen) at 1:1000. Ovaries were mounted in Vectashield with DAPI (Vector Labs). The cell membranes were labelled with CellMask™ Orange Plasma Membrane Stain or CellMask™ Deep Red Plasma Membrane Stain (ThermoFisher Scientific).

### Immunostainings

Ovaries from adult flies or imaginal wing discs from third instar larvae were dissected in phosphate-buffered saline (PBS) and fixed for 20 min in 4% paraformaldehyde and 0.2% Tween in PBS. The tissues were then incubated in 10% bovine serum albumin (in PBS) to block for one hour at room temperature. The incubation with primary antibody was performed at 4C overnight in PBS, 0.2% Tween and 1%BSA.

Immunostaining on pupal eye disc was performed as described in (Richard et al., 2006).

### Imaging

Fixed preparations were imaged using an Olympus IX81 (40×/1.3UPlan FLN Oil or 60×/1.35 UPlanSApo Oil). Live imaging was performed using a Leica SP8 (63×/1.4 HCX PL Apo CS Oil) or Olympus IX81 (40×/1.3 UPlan FLNOil or 60×/1.35 UPlanSApo Oil) inverted confocal microscope. For live observations, ovaries were dissected and imaged in 10S Voltalef oil (VWR Chemicals).

### Drosophila genetics

Follicle cell clones of *imp*, *Dl*, *bel*, *dcr-1* and *kuz* were induced by incubating larvae or pupae at 37°C for two hours every twelve hours over a period of at least three days. Adult females were dissected at least two days after the last heat shock. Wing imaginal disc clones and eye imaginal disc clones of *imp* were induced by heat shocking first and second instar larvae for 30min per day over a period of 2 days. Larvae were dissected at least one day after the last heat shock.

## Acknowledgements

We thank Sarah Bray, Francois Schweisguth, Wu-Min Deng, Alain Debec and Andrea Brand and their labs for fly stocks and Akira Nakamura for antibodies; the Developmental Studies Hybridoma Bank, Kyoto Stock Center (DGRC) and Bloomington Drosophila Stock Center (BDSC) for antibodies and fly stocks; D. St J. lab for technical assistance, helpful comments and criticism.

## Author contributions

W.F., C.F. and D. St J. designed the study. W.F and C.F performed all experiments. W.F. and D. St J. prepared the manuscript.

---

**Supplementary Fig 1.**
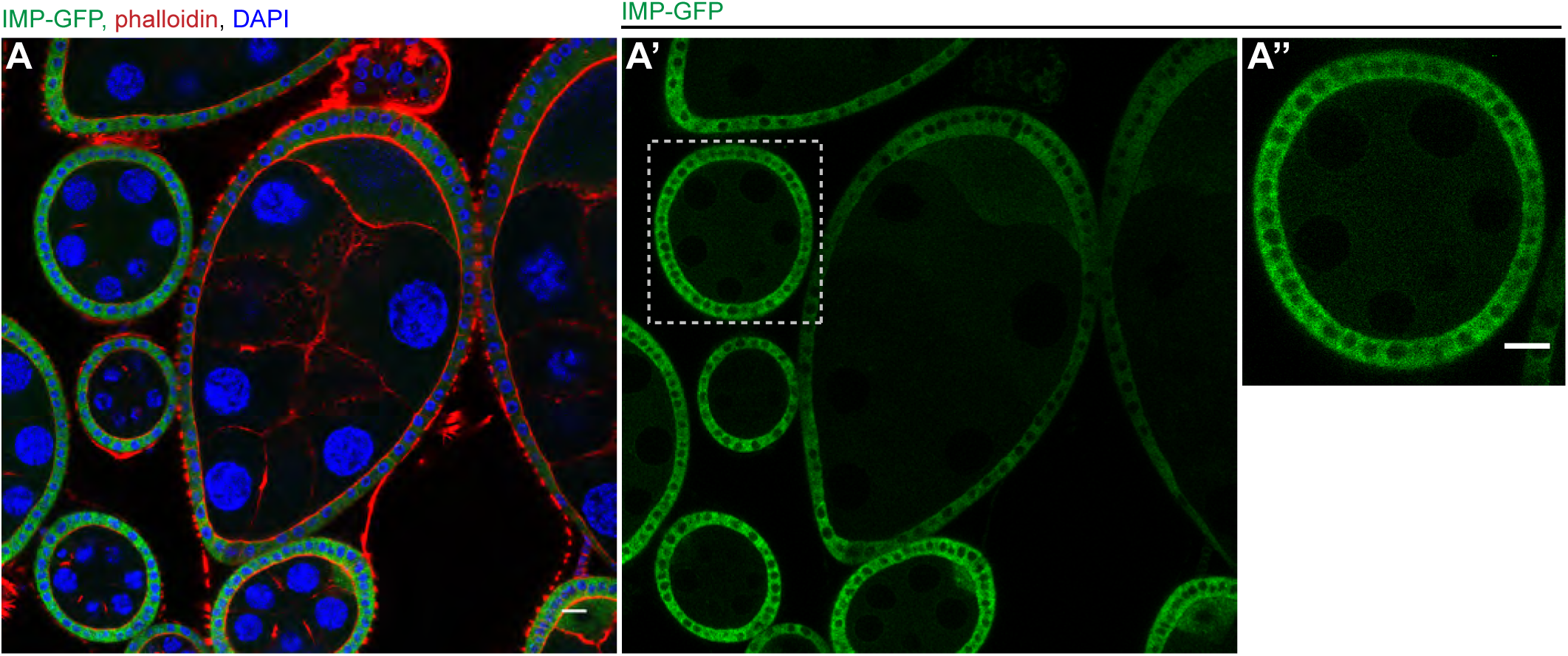
IMP is uniformly distributed throughout the cytoplasm of the follicle cells. (A-A”) Wild-type egg chambers expressing IMP-GFP (green) from a protein trap insertion in the first intron, stained for actin (red) and DNA (blue). (A”) shows the uniform distribution of IMP in the follicle cells at stage 6 when Dl/N signalling occurs.

**Supplementary Fig. 2.**
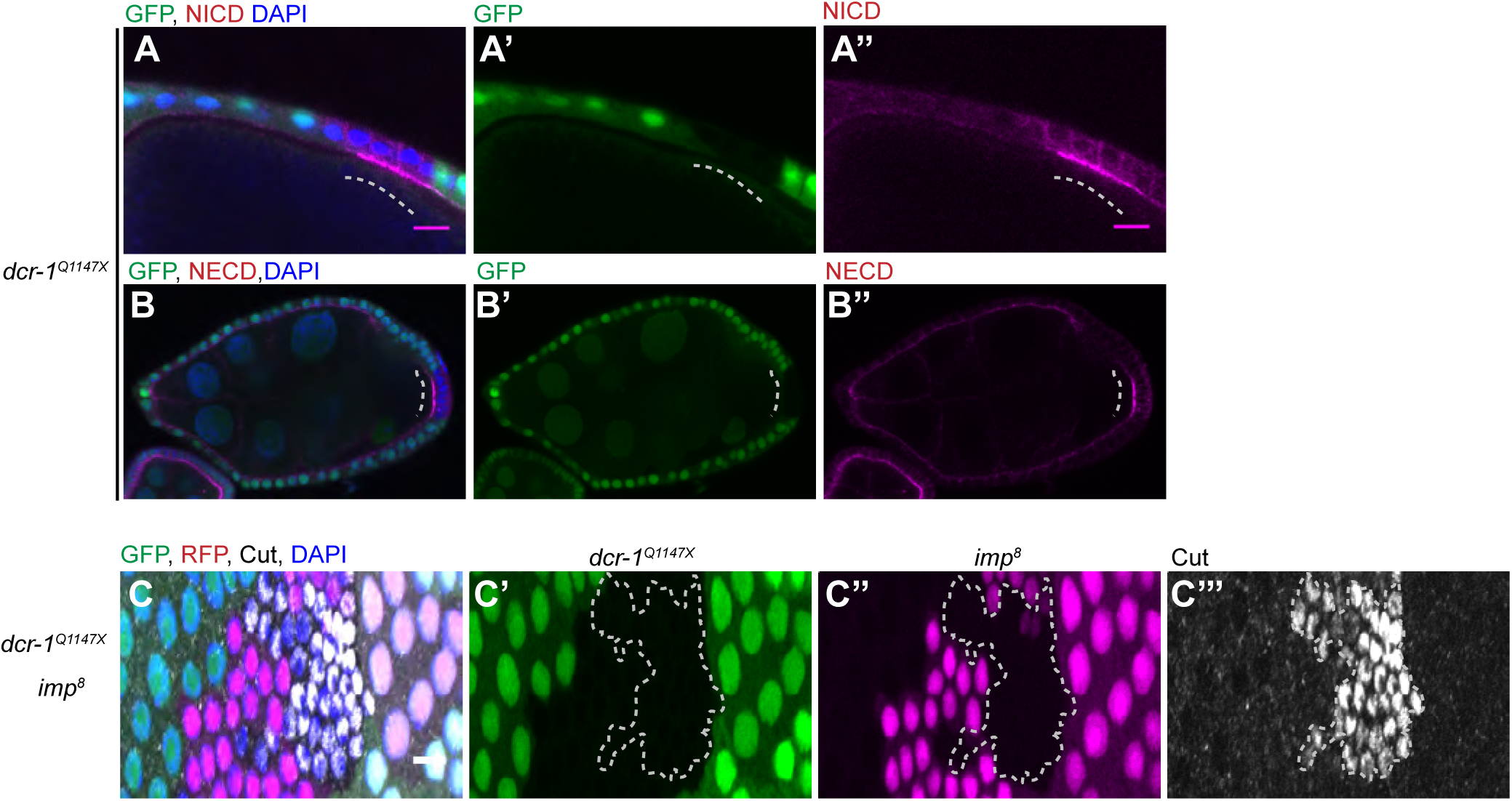
IMP does not act through the micro RNA pathway. (A-B) Stage 7 egg chambers containing *dcr-1*^Q1147X^ mutant clones marked by the loss of GFP (green). The mutant cells retain high levels of NICD (magenta in A, A’, A”) and NECD (magenta in B, B’, B”) at the apical membrane. (C-C”’) A stage 9 egg chamber containing both *imp^8^* mutant clones marked by the loss of RFP (magenta) and *dcr-1*^Q1147X^ mutant clones marked by the loss of GFP (green), stained for Cut (white). Cut is still expressed in the double mutant cells (surrounded by the dashed line), but not in the single mutant cells, indicating that loss of Dicer and IMP has an additive effect on the delay to Notch signalling.

**Supplementary Fig. 3.**
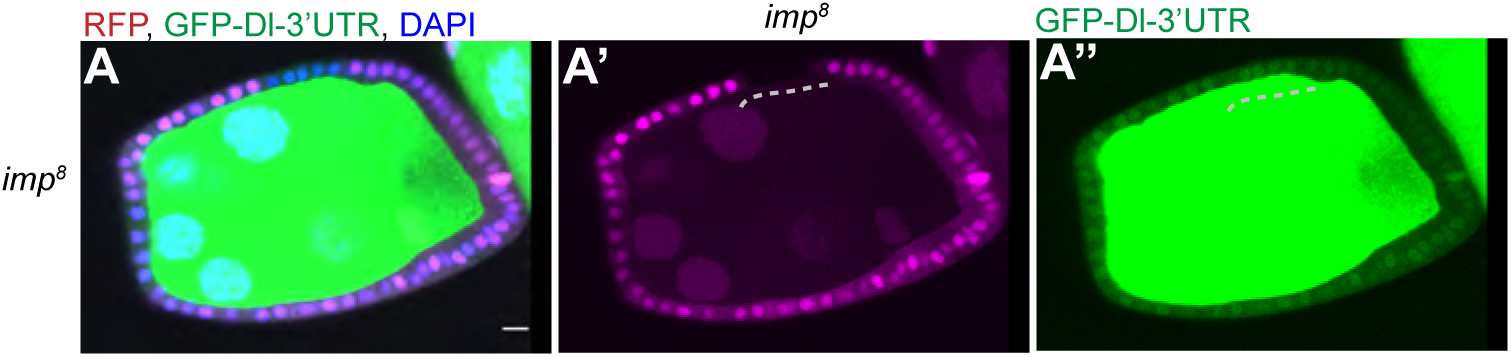
IMP does not control the expression of a GFP-Dl 3’UTR sensor. (A, A’, A”) *imp*^8^ mutant cells marked by the loss of RFP (magenta) express the same levels of GFP from the GFP-Dl-3’UTR sensor as wild type cells.

